# RC-GNN: A predictive model of enzyme-reaction pairs

**DOI:** 10.1101/2025.06.22.660952

**Authors:** Stefan C. Pate, Eric H. Wang, Linda J. Broadbelt, Keith E.J. Tyo

## Abstract

Uncharacterized functions of enzymes represent untapped opportunity to develop therapeutics, unlock the sustainable synthesis of materials, and understand the evolution of life-sustaining metabolic networks. Uncharacterized enzymes and reactions, generated by protein language models and computer-aided synthesis tools, respectively, make up a large part of this opportunity. Given the technical complexity of high-throughput enzymatic activity screens, predictive models are needed that can pre-screen enzyme-reaction pairs *in silico*. We present Reaction-Center Graph Neural Network, (RC-GNN) a model capable of predicting whether an enzyme, represented by an amino acid sequence, can significantly catalyze a given reaction, represented by its full set of reactants and products. We explicitly evaluated RC-GNN’s generalization to queries highly dissimilar from those present in the training dataset. In the most difficult conditions tested, our models achieve 0.88 and 0.84 ROC-AUC on classification tasks featuring globally selected and synthetic negatives, respectively. On a time-based split an RC-GNN achieved 0.91 ROC-AUC. The ability to successfully make predictions on enzymes and reactions distinct from those used during training makes RC-GNN especially useful for both metabolic engineers and evolutionary biologists who need to reason about uncharacterized enzymatic reactions.

## Introduction

A complete mapping of enzyme sequences to the reactions they catalyze would enable progress across multiple fields: pharmaceutics, biomanufacturing^1,2^, physiology, and evolutionary biology^3–7^. Over decades, researchers have characterized the function of thousands of enzymes, namely the canonical reactions that they catalyze. Nevertheless, it is widely believed that our collective knowledge is far from complete. In large part, the missing pieces are secondary functions, also known as promiscuous activities of enzymes, or the underground metabolism^8^. Typically, these activities are less efficient than the primary function but far more efficient than the uncatalyzed reaction. To an evolutionary biologist, they represent both vestiges of evolutionary history and sources of evolvability required for future phenotypic change. To a biotechnologist, they represent both a challenge and an opportunity. Secondary functions required for a synthesis pathway can be elicited with protein engineering. On the other hand, they are sources of unanticipated adaptations to a metabolic engineer’s purposeful genetic perturbation.

To put the experimental challenges associated with mapping enzyme-reaction pairs into context, testing all possible enzyme-reaction pairs from the dataset studied in this work would require hundreds of millions of experiments. Furthermore, this does not include novel, predicted enzymes and reactions, as one considers in computer-aided biosynthesis. At the same time, throughput of a systematic screen of enzymatic functions is limited by a lack of detection methods applicable to such a wide range of reactants and products^8^.

Given this challenge, a critical question is what experiments should be prioritized in order to most efficiently increase our knowledge of enzymatic function. While theoretical^9^ and heuristic understanding of enzymatic reaction mechanisms can greatly constrain this search, the range of structures (of both substrates and enzymes) and functions is great, and the relationship between them is rich. Machine learning (ML) is an appropriate tool to learn such a complicated relationship from data. An ML model can inform decisions on experiment prioritization, and its parameters can be updated with the experimental results to close the loop of semi-automated exploration.

Such a model must accept two inputs, an enzyme and a reaction, and score the hypothesis that the former catalyzes the latter. Previous work has formulated a variety of related tasks to the one just described. One group of models outputs Enzyme Commission (EC) numbers given an amino acid sequence or reaction^10–16^. Predicting EC numbers is useful but limited in that it is not a precise description of enzymatic function^17–19^. A second group of tools looks up enzymes empirically associated with characterized reactions similar to a query reaction or substrate^20–23^.

These models are limited to the task of finding enzymes already associated with known reactions and mostly utilize non-differentiable methods to perform the lookup. A third group of models scores the interaction between the enzyme and a putative substrate^24–26^. While it is useful to know if a substrate and enzyme interact, this fact alone leaves ambiguity around the transformation that the substrate undergoes. For example, pyruvate may undergo several reactions including reduction to lactate (EC: 1.1.1.28), decarboxylation to acetaldehyde and carbon dioxide (EC: 4.1.1.1), or carboligation to acetolactate (EC: 2.2.1.6). A primary task of computer-aided biosynthesis is to predict an enzyme to catalyze a reaction^2^, and the design of an experiment to test such a prediction includes at a minimum an amino acid sequence and the chemical structures of all reactants and products. Recent works have moved toward this more precise formulation of the enzyme function prediction task^27,28^.

Here we build on earlier enzyme function prediction work in several ways. We curate a dataset of high-fidelity enzyme-reaction pairs supported by direct experimental evidence only. Based on these underlying positive pairs, we develop a battery of model evaluations involving several approaches to data splitting and negative sampling. Finally, we develop a series of models which show improved performance on enzyme-reaction pair prediction over previous studies. In their Graph Neural Network (GNN) reaction encoders, the best-performing of these models use a chemistry-inspired form of graph augmentation based on the reaction center (RC). RC information has been used successfully as data features^29^ and labels^30^ on other tasks in cheminformatics such as reaction classification and retrosynthesis. The RC-GNN models developed here generalize well to reactions highly dissimilar to those in the training set as well as prospectively on a time-based split. Additionally, the RC-GNN models are both parameter and sample efficient, and, when they err, do so in a chemically reasonable manner, positioning them well for practical use.

## Results

### Preparing a dataset of enzyme-reaction pairs

We acquired experimentally validated enzyme-reaction pairs from UniProt, which features cross-referenced enzymatic reactions from the Rhea database^31,32^. We note that enzymatic reaction databases like Rhea are likely biased toward primary, native functions over promiscuous ones.

However, there is appreciable multiplicity in both the enzyme-to-reaction and reaction-to-enzyme mappings (Fig. S2). In contrast to previous data curation efforts^28,33^, we restricted our dataset to manually-annotated pairs with direct evidence, not inferred from homology, and removed subunits of multimeric enzyme complexes. A lack of true negative samples is a common issue in binary classification tasks and was true in our case. In lieu of true negatives, practitioners use one of several strategies to sample negative data points^34,35^. Two common approaches are global selection from all unobserved pairs, and the use of an auxiliary similarity metric to bias negative samples to be more or less similar to observed pairs^34^. Prior work on enzyme function prediction has used auxiliary similarity metrics to bias negative samples in both directions^24,25^. Given this challenge, we took a two-pronged approach to negative sampling. In the first, we employ global selection of negatives, relying only on a basic assumption that the underlying adjacency matrix of enzymes and reactions is sparse, and avoiding any assumptions about the structure-function relationship, which can exhibit discontinuities as discussed in other domains of cheminformatics^36^. The adjacency matrix of the observed enzyme-reaction pairs has a density of 0.03%. The density of the true adjacency matrix being inferred is unknown^8^; however, assuming current knowledge covers at least 10% of enzyme promiscuity, the probability of misassigning true positives is less than 3%. While global sampling has the advantage of avoiding misassignment of true positives, its limitation is its bias toward relatively “easy” negatives, i.e., enzymes matched with reactions which are highly dissimilar to those they have been characterized to catalyze. Therefore, we conducted experiments with negative pairs sampled using the alternate reaction center hypothesis (ARC)^37^. This approach generates negative data by applying minimal transformations^18^ to moieties which are identical to those acted on in observed reactions, but which are surrounded by different structural context. This results in “harder” negative samples, those which are quite similar to observed reactions. See the methods section for a detailed explanation of ARC and Figure 4 for an illustration. Summary statistics for the resulting dataset are reported in Table 1, as well as in Figures S2 and S3.

**Table 1.** Summary dataset statistics. ARC = alternate reaction center.

|  | Global negative sampling | ARC negative sampling |
| --- | --- | --- |
| Number of enzymes | 24,523 | 24,523 |
| Number of reactions | 6,460 | 76,292 |
| Number of positive pairs | 49,071 | 49,071 |

A major focus of this work is to demonstrate model generalization to enzymes and reactions highly dissimilar to the training set. To this end, we utilized protein and reaction similarity metrics to precisely quantify (dis)similarity. This approach has been used in prior ML tasks involving proteins to avoid overly optimistic estimates of generalization error^38^. To address this issue on the reaction side, we developed reaction center maximum common subgraph (RCMCS). Analogously to global sequence identity (GSI), which we use to measure pairwise protein similarity, RCMCS is based on an alignment of reaction graphs. These similarity metrics were used in a stratified similarity split procedure to ensure that models were evaluated on test data points spanning a range of dissimilarity from the training set. See the methods section for details.

As a demonstration of the validity of RCMCS as an enzymatic reaction similarity score, we classified reactions with EC number in a leave-one-out cross validation procedure (Table 2). For each query reaction, EC numbers were predicted by retrieving the *k* most similar reactions from the data set. RCMCS similarity more accurately retrieved EC numbers than did Tanimoto similarity of DRFP vectors, an optimization-free reaction fingerprinting procedure^39^.

**Table 2.** RCMCS similarity aligns with EC number classification. Comparison of RCMCS and DRFP similarity in leave-one-out EC number prediction.

|  | Top-1 |  | Top-3 |  | Top-5 |  | Top-10 |  |
| --- | --- | --- | --- | --- | --- | --- | --- | --- |
|  | RCMCS | DRFP | RCMCS | DRFP | RCMCS | DRFP | RCMCS | DRFP |
| EC level 1 | 0.970 | 0.893 | 0.980 | 0.947 | 0.985 | 0.963 | 0.993 | 0.978 |
| EC level 2 | 0.949 | 0.807 | 0.961 | 0.874 | 0.965 | 0.900 | 0.974 | 0.929 |
| EC level 3 | 0.924 | 0.786 | 0.946 | 0.837 | 0.950 | 0.867 | 0.958 | 0.899 |

### Constructing a model to map enzymes and reactions into a joint embedding space

To predict the catalysis of a reaction by an enzyme we took the approach of learning representations of both. We develop models that map each item in the pair to the same vector space. With this, we can score whether the enzyme in a pair would catalyze the reaction by taking a dot product between the two items’ vector embeddings. This architecture of model requires two encoders (Fig. 1a). The protein encoder consists of the ESM-1b transformer model which maps an amino acid sequence to a 1280-dimensional vector^40^. The ESM vector embedding is then linearly transformed with a matrix learned from the current task. The architecture of the reaction encoder, which maps the SMILES^41^ representation of a reaction to a vector embedding, is a major focus of the present work.

**Figure 1.**
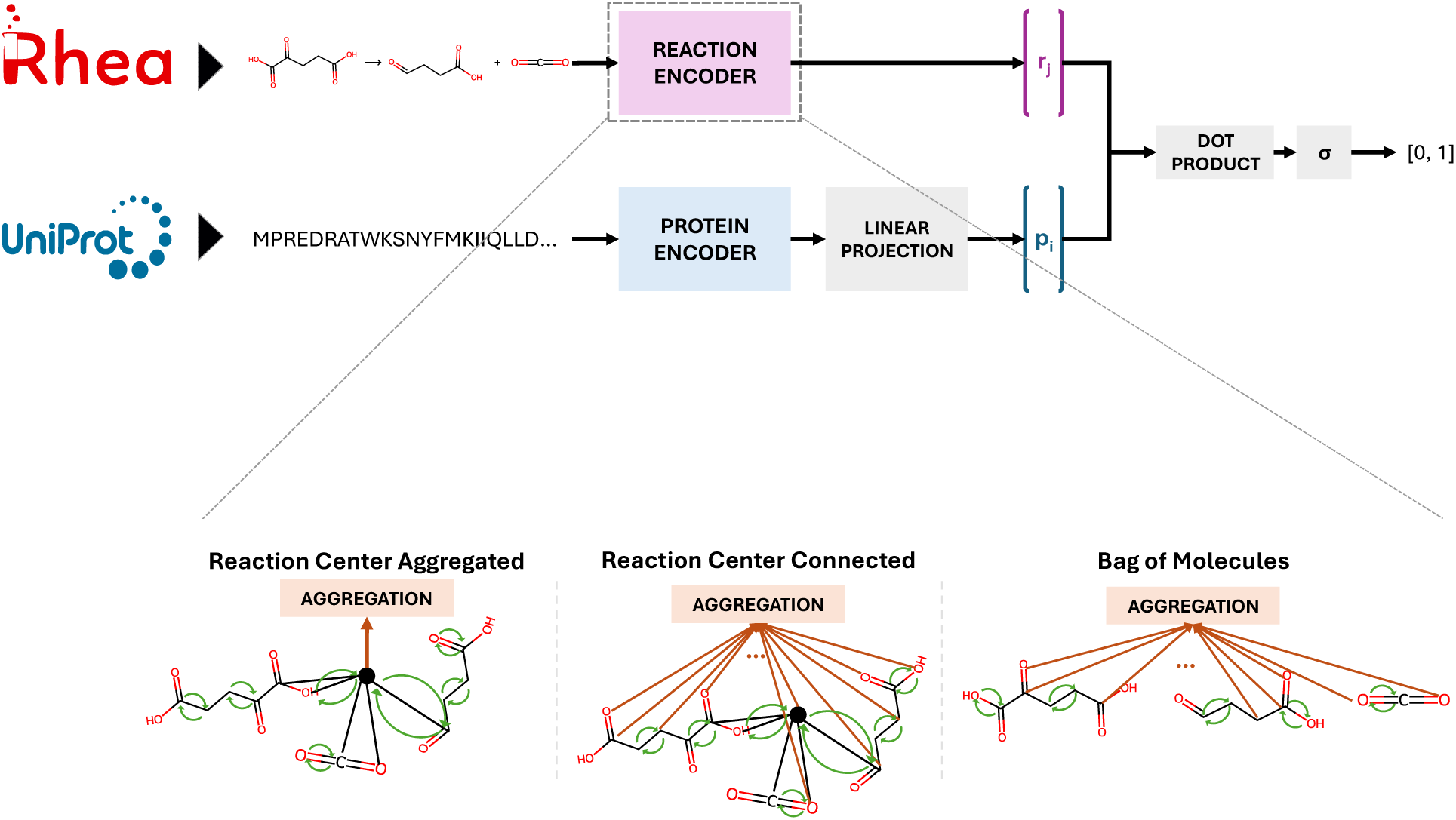
Overview of the model architecture. (Top) All models take as input a pair of reaction, encoded as a SMILES string, and protein, encoded as an amino acid sequence. Each item in the pair is transformed into a vector representation by an encoder. Finally, the pair of vectors is converted into a score, which is a function of their dot product. (Bottom) Several variants of reaction encoders were compared. Bag of molecules (right) represents a reaction as the mean of all atom embeddings following several rounds of message passing. RC aggregation (left) represents a reaction with the embedding of a virtual node which has been connected to RC atoms via virtual edges. RC connected (center) also introduces a virtual node connected to the RC but represents a reaction as the mean of all atom embeddings as in bag of molecules.

GNNs have been used to generate learned molecular embeddings^42^, and the Chemprop package offers a suite of GNN functionality bespoke to cheminformatics^43^. Building on this prior work, we developed and compared three GNN-based reaction encoders (Fig. 1b and Table 3). The first, bag of molecules, represents a reaction with the mean embedding of atoms across all reactants and products, each of which is generated after a number of message passing steps. The RC aggregated encoder augments the graph of reactants and products with a virtual node as done previously in molecular property prediction^44^. In our case, we augmented the graph by connecting a virtual node to the RC atoms by virtual edges. Though initially stateless, the virtual node inherits a combination of atom and bond features through message passing. This combination includes features within *k* bonds of the RC where *k* is one less than the number of message passings. The RC connected encoder is a hybrid between the former two. Like RC aggregated, it augments the reaction graph with a virtual node, but the reaction is represented as the mean atom embedding like in the bag of molecules encoder.

**Table 3.**
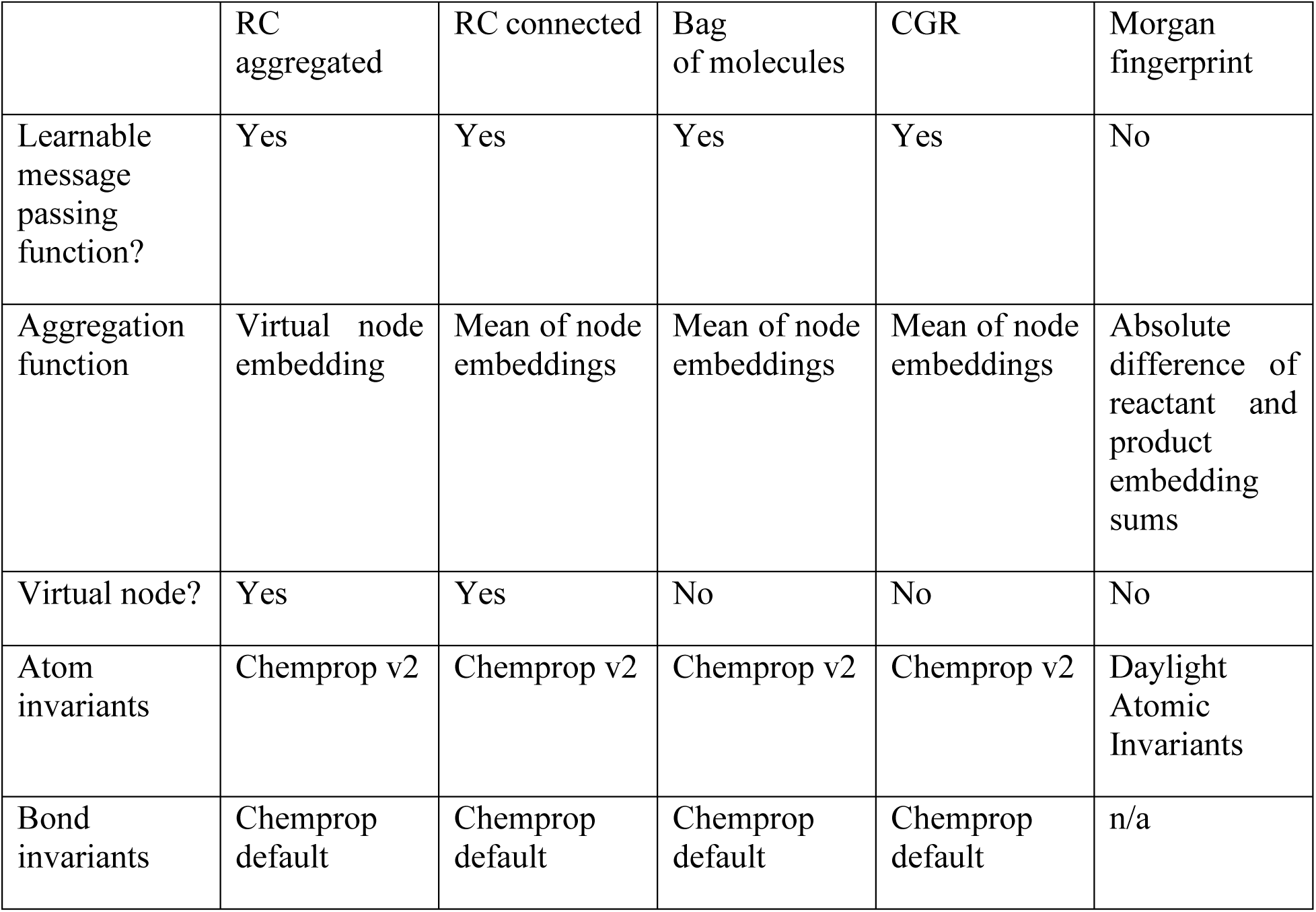
Key differences among reaction encoders. RC = reaction center; CGR = condensed graph of reaction.

Additionally, we compared these three GNN encoders with a condensed graph of reaction (CGR) encoder^45,46^ implemented in Chemprop, and a linear combination of reactant and product extended-connectivity fingerprints (ECFP)^47^ also referred to as Morgan fingerprints. For a summary of reaction encoders, see Table 3. Like the GNNs, the Morgan fingerprint encoder works by iterative message passing, but without a learnable weight matrix combining the messages. On each side of the reaction, substrate fingerprints were summed. Then the absolute difference between the summed fingerprints of each side was taken as the reaction fingerprint.

Finally, we benchmarked the suite of reaction encoders described above to several encoders and fingerprints previously described in literature. These include a GNN-based pseudo-transition-state encoder which we refer to as CLIPZyme^27^, an ECFP-based reaction fingerprint, DRFP^39^, and a transformer-based vector embedding which was fine-tuned on USPTO reaction classification, RXNFP^27,39,48^. These latter two fingerprint vectors were transformed by learned matrix or multi-layer perceptron (Table S1). In all evaluation experiments, benchmark reaction encoders were trained jointly with the linear protein encoder described above. Note however, that in the time-based split, we utilize the previously trained CLIPZyme model which consists of an Equivariant GNN protein encoder (EGNN)^27^. We describe this experiment further in the section, “Models perform well prospectively on time-based split”.

### Successfully predicting unseen enzyme-reaction pairs

We estimated generalization error on test sets generated using a stratified similarity split technique. This splitting technique ensures that the maximum pairwise similarity of a test data point to train data points varies over a wide range. In other words, the test set consists of a subset of data points with low similarity to the training set, a subset with high similarity to the training set, and so on for several intermediate levels of similarity. Since our data points are pairs of enzymes and reactions, we carry out the stratified similarity splitting procedure twice, using RCMCS and GSI for reactions and enzymes, respectively. See Methods for details on stratified similarity split. See Fig. S1 for the distribution of pairwise similarities over the whole dataset and in our test set, relative to the training set.

We find in general that splitting based on pairwise reaction similarity, RCMCS, leads to a more challenging task than GSI-based splitting. While all models perform above chance at labeling unseen enzyme-reaction pairs as positive or negative, the GNN and RXNFP models outperform the reaction fingerprints Morgan and DRFP on reaction-based stratified similarity splits (Fig. 2a and 2b). The RC connected model achieved the highest ROC-AUC score of 0.940, followed by RC aggregated (0.938) (Fig. 2b). All GNN-based and Morgan fingerprint models performed comparably on the GSI-based split (Fig. 2c and 2d). Such robustness to protein dissimilarity has been observed before^24^.

**Figure 2.**
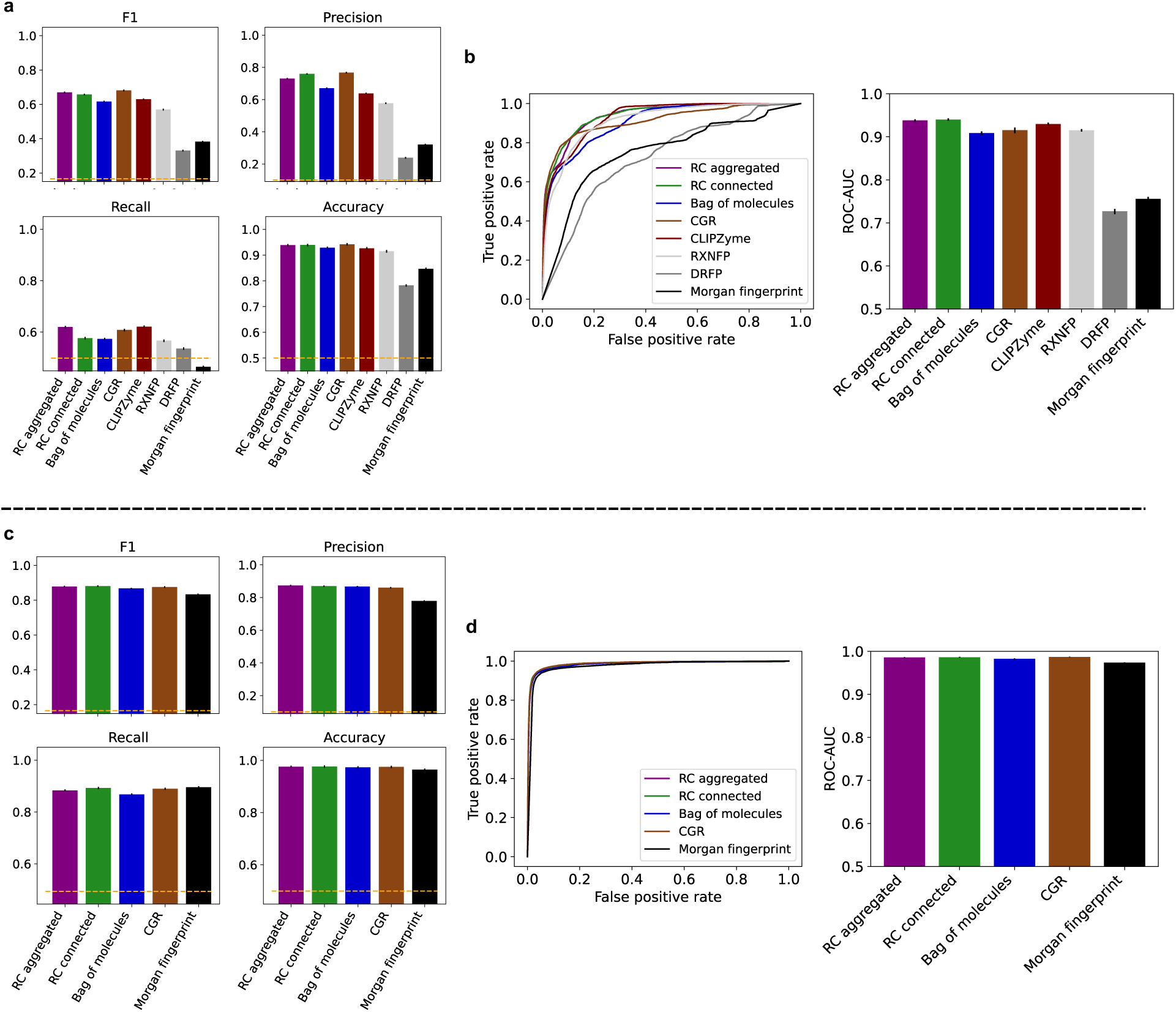
Models successfully predict enzyme-reaction pairs. (a) Classification performance metrics plotted for each type of model. (b) Left, ROC curves plotted for each type of model using test data. Right, ROC-AUC scores. In both (a) and (b), stratified similarity split with RCMCS as the similarity metric was used to split data. (c) and (d) Same as (a) and (b), respectively, but using GSI as the similarity metric in stratified similarity split. Chance-level performance is given by the orange dashed lines in (a) and (c). Error bars are 95% confidence intervals computed by bootstrapping (n=20).

### Chemistry-inspired graph augmentation leads to robust model performance

We sought to understand the relationship between reaction similarity and the difficulty of predicting enzyme function by leveraging our stratified similarity splitting technique. We hypothesized that a model would perform worse where query reactions look quite different from the training set. This is consistent with the general notion that machine learning models excel at interpolation within a distribution of data but not extrapolation beyond it^49^. Alternatively, it could be true that the task becomes difficult in the other extreme where query reactions are quite similar, but with subtle differences that affect function to an outsized extent^36^.

By separating our test data points into subsets based on their similarity to the training set, we support the first hypothesis that dissimilar data points make more challenging queries (Fig. 3). Nonetheless, all models maintain classification performance appreciably above chance over a substantial range of reaction similarity. The most pronounced drop in performance occurs in the [0.0, 0.4) RCMCS similarity tranche where, overall, GNN and RXNFP models outperform the other reaction fingerprints. Moreover, the RC connected model distinguishes itself from the rest of the GNNs on this most difficult subset of data, maintaining 0.88 ROC-AUC (Fig. 3a). The second best performing model in this difficult subtask was RC aggregated in terms of decision-threshold-invariant measures (ROC-AUC = 0.87); however, the CGR architecture outperformed both on F1, for which an optimal decision threshold was selected using validation data (Fig. 3b). As an illustration of our reaction similarity metric, RCMCS, we show three reaction pairs with varying levels of similarity shown on the left of the chemical equations (Fig. 3c). We highlight in green the maximum common subgraph inclusive of the RC.

**Figure 3.**
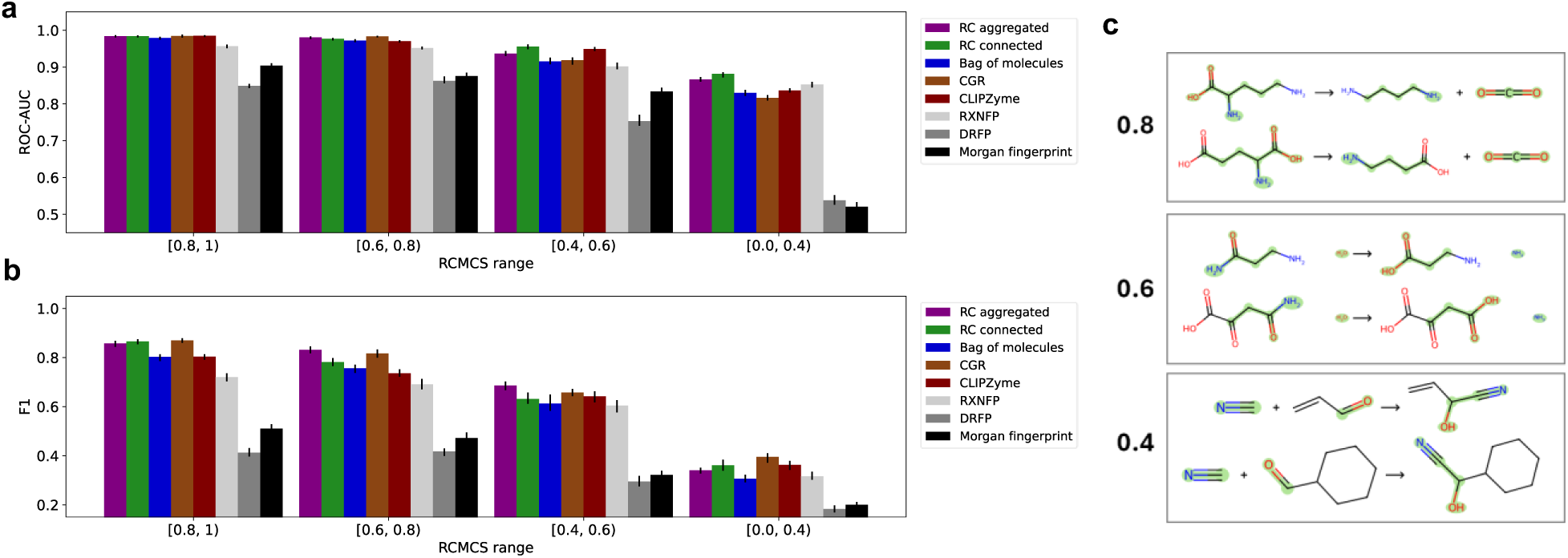
Model performance is robust over a range of reaction similarity. (a) Test set F1 score plotted against maximum RCMCS similarity between test data points and train data points. (b) Same as (a) for F1. (c) Illustrative examples of RCMCS similarity scores for three pairs of reactions. The maximum common subgraph inclusive of the RC is highlighted in green. Error bars are 95% confidence intervals computed by bootstrapping (n=20).

We repeated this analysis with a protein-based similarity metric, GSI. All models generalized extremely well to even the most dissimilar subset of enzymes (Fig. S4). This suggests that generalization to highly dissimilar reactions is more challenging than generalization to highly dissimilar enzymes. Although we were motivated to find the point at which performance would decline, we were limited to a lower bound of 30% GSI beyond which the dataset collapsed into a single cluster, precluding stratified similarity splitting.

### Models can distinguish observed pairs from hard, ARC-sampled negatives

Similarity splitting, and RCMCS-based splitting in particular, challenges models to make inferences far from regions of chemical space represented by their training data. We next challenged models to distinguish between observed enzyme-reaction pairs, and pairs which look very similar to those observed, but which we can infer are likely to be negative samples. We illustrate the ARC-based negative sampling approach with the case of isoleucine hydroxylation (EC 1.14.11) (Fig. 4). In the curated dataset, there are hydroxylations of isoleucine at two different positions, catalyzed by two different enzymes. As we have filtered our data to include only enzyme-reaction pairs with direct experimental evidence, we assume the lack of reporting of a reaction, equivalent in its reactants and reaction center to a reaction reported as catalyzed by a certain enzyme, strongly implies its infeasibility. Cases (i) and (ii) depict examples where we infer negative enzyme-reaction pairs from this lack of reporting, though the reactions of the inferred negative pairs have been reported catalyzed by a different enzyme in our dataset (Fig. 4). Case (iii) represents another situation in which an isoleucine hydroxylation occurs at a position never reported. We infer in this case one negative pair per enzyme known to catalyze isoleucine hydroxylation at other positions (Fig. 4).

**Figure 4.**
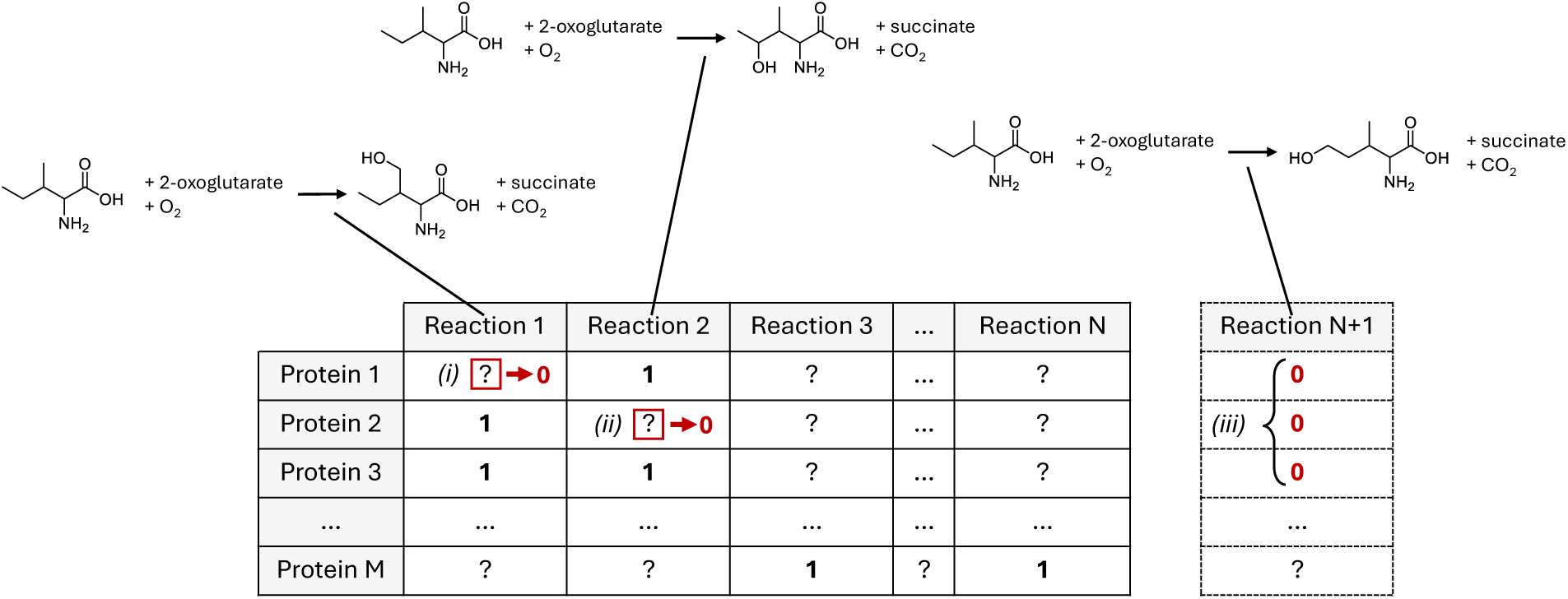
Illustration of negative sampling via alternate reaction center hypothesis. Observed reactions 1 and 2 have equivalent reactants and are described by equivalent minimal reaction rules, yet hydroxylation occurs on different carbons, yielding different products. (i) It can be inferred that protein 1 cannot catalyze reaction 1 since it did not transform the displayed reactants into the products of reaction 2 in experiment. (ii) It can be inferred that protein 2 cannot catalyze reaction 2 for the same reason as (i). (iii) It can be inferred that none of proteins 1-3 can catalyze unobserved reaction N+1 for the same reason as (i) and (ii).

On top of ARC-negative sampling, we layer reaction-based data splitting of two extremes. In the easiest extreme, unique reactions, along with all their positive and negative enzyme pairs, are randomly held out as test data. Models performed comparably to the case of global selection of negatives in the most forgiving range of RCMCS, [0.8, 1.0) (Fig. 5 a & b). This motivated us to push data-splitting to the opposite extreme, randomly selecting unique reaction centers (equivalently minimal reaction rules) to hold out as test data. This is equivalent to imposing an RCMCS bound of 0.0 (inclusive). Combined with ARC-based negative sampling, this produced the most difficult task of our study. The RC aggregated and RC connected models led performance in this task with ROC-AUC of 0.84 and 0.82, respectively (Fig. 5d). The CGR model led in F1 score (0.60), followed by RC connected (0.50). We observed opposite tradeoffs between precision and recall, with CGR, RC connected, and RC aggregated exhibiting high precision but poor recall. Conversely, CLIPZyme and DRFP models exhibited higher recall than precision. These were the results for the decision thresholds selected through cross validation to maximize F1. Practitioners are advised to select the threshold most appropriate for their goals.

**Figure 5.**
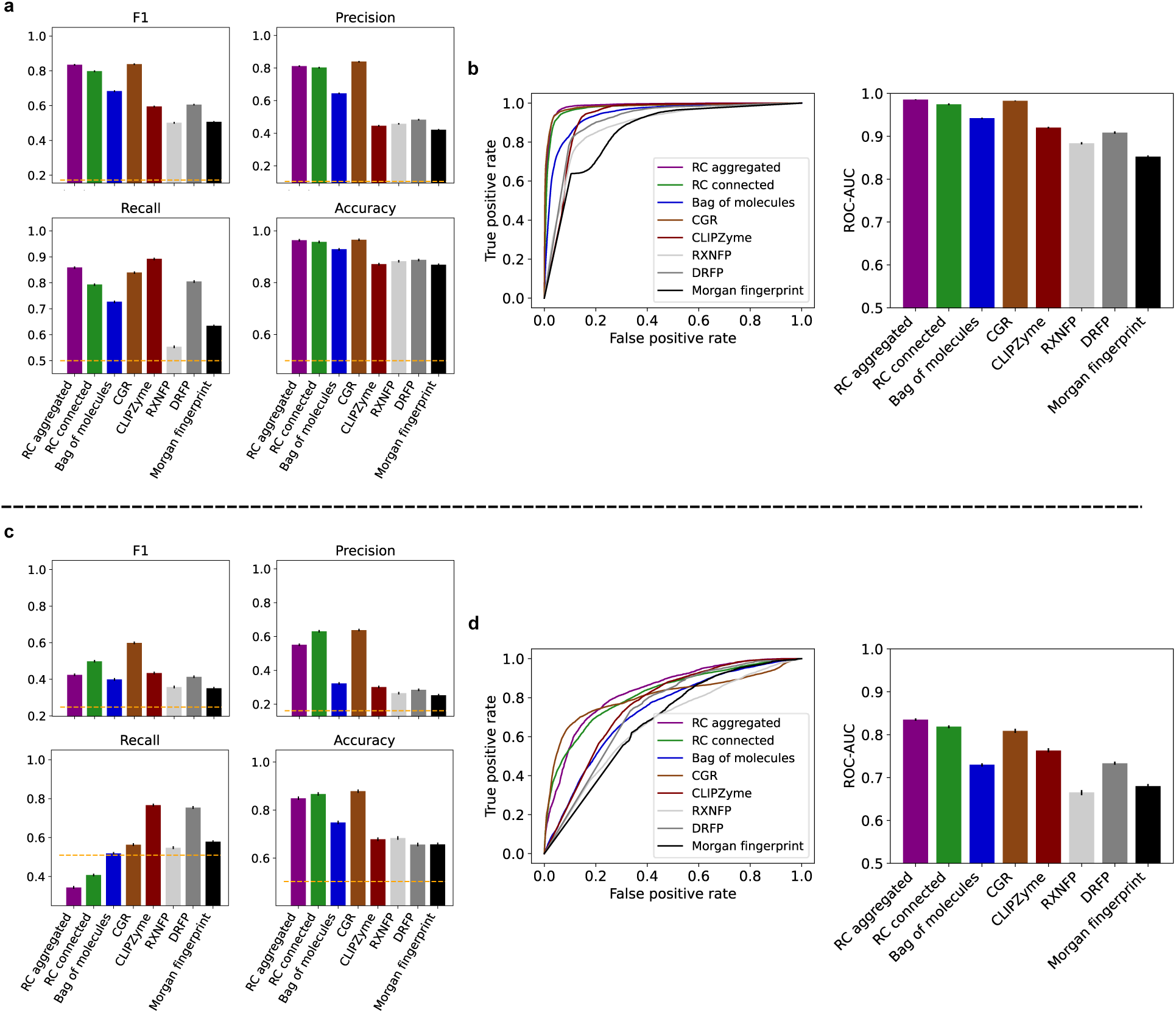
Models successfully distinguish between observed positive and ARC-negative pairs. (a) Classification performance metrics plotted for each type of model. (b) Left, ROC curves plotted for each type of model using test data. Right, ROC-AUC scores. In both (a) and (b), negatives were sampled using the ARC hypothesis and data were split by random selection of reactions, equivalent to an RCMCS bound of 1 (exclusive). (c) and (d) Same as (a) and (b), respectively, but data were split by random selection of reaction centers, equivalent to an RCMCS bound of 0 (inclusive). Chance-level performance is given by the orange dashed lines in (a) and (c). Error bars are 95% confidence intervals computed by bootstrapping (n=20).

### Models perform well prospectively on time-based split

To complement our above-described evaluations, we performed a time-based split to evaluate models’ ability to make prospective predictions on new literature (Table 4). We additionally use this task to benchmark against the full workflows of CLIPZyme – inclusive of their data set, contrastive training objective, optimized pseudo-transition-state reaction encoder, and EGNN-based protein encoder – and ReactZyme. Our time-based test split contains enzymatic reactions from across all EC classes and is challenging in the sense of having a distinct composition of reaction centers from the training data set (Fig. S8). On the full set of enzyme-reaction pairs, the bag of molecules model showed the best performance, followed closely by RC connected. RC aggregated trailed these other two models by more than in other tasks, suggesting that information from outside of the peri-reaction-center subgraph was useful. To make a fair comparison, we calculated ROC-AUC on a subset of the data since CLIPZyme rejects 64 out of 371 enzymes for having sequences which are longer than 650 residues^27^. On this subset, the RC connected model performed best, followed by CLIPZyme and bag of molecules. RC connected and bag of molecules achieved such performance using nearly four orders of magnitude fewer trainable parameters than CLIPZyme, with the majority of this difference stemming from the latter model’s EGNN-based protein encoder. The models of this study were also more sample efficient. Though the number of positive enzyme-reaction pairs in our data set is similar to that used by CLIPZyme, the current work samples three orders of magnitude fewer negative pairs.

**Table 4.** Performance on time-based split. ROC-AUC is reported both for the full time-based split and for the subset for which CLIPZyme is able to make predictions.

|  | ROC-AUC | ROC-AUC<br>filtered | Number of<br>trainable<br>parameters | Number of<br>negative training<br>samples |
| --- | --- | --- | --- | --- |
| RC aggregated | 0.815 | 0.825 | $2.5 \times 10^5$ | $2.0 \times 10^5$ |
| RC connected | 0.887 | **0.905** | $7.3 \times 10^4$ | $2.0 \times 10^5$ |
| Bag of molecules | **0.889** | 0.888 | $3.5 \times 10^4$ | $2.0 \times 10^5$ |
| CGR | 0.816 | 0.848 | $5.7 \times 10^5$ | $2.9 \times 10^5$ |
| CLIPZyme | n/a | 0.893 | $2.3 \times 10^8$ | $2.1 \times 10^8$ |
| ReactZyme | 0.733 | 0.743 | $1.3 \times 10^6$ | $1.6 \times 10^6$ |
| RXNFP | 0.859 | 0.858 | $8.5 \times 10^4$ | $1.5 \times 10^5$ |
| DRFP | 0.762 | 0.783 | $8.6 \times 10^4$ | $1.5 \times 10^5$ |
| Morgan<br>fingerprint | 0.754 | 0.806 | $2.0 \times 10^5$ | $1.5 \times 10^5$ |

### RC aggregated model learns chemically meaningful information

Although it was not the best performing model in every experiment, the RC aggregated model was consistently competitive with other baseline models. This was surprising given it is strongly constrained to utilize information from within few bonds of the RC. We wondered whether this constraint had the effect of focusing the model on chemically meaningful information. To test this hypothesis, we analyzed false positive errors made by the evaluated models. We compared the reaction that was wrongly predicted to all reactions empirically shown to be catalyzed by the query enzyme. Compared to other models, the false positive errors of RC aggregated tended to be more similar to the ground truth reactions as measured by number of matching EC digits (Fig. 6 a and b). For example, 47.64% of RC aggregated false positives shared two or more EC digits with true positive reactions. CGR consistently showed the next highest values on this analysis with 41.25%. In contrast, bag of molecules and DRFP made false positive errors farther afield from ground truth, scoring 29.08% and 11.22%, respectively. The same trends were observed using the RCMCS similarity score instead of EC number matches (Fig. S9).

**Figure 6.**
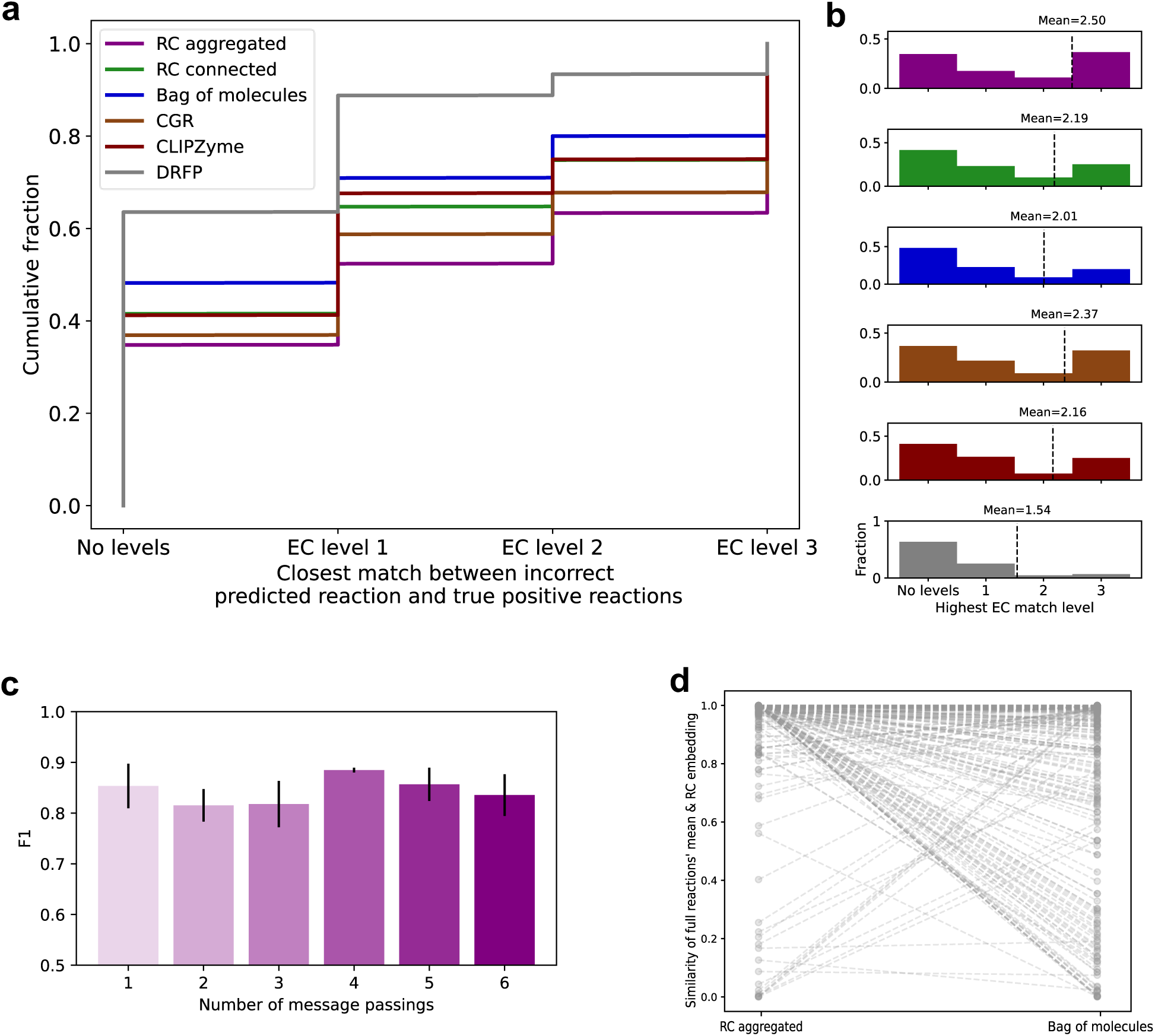
RC aggregated learns chemically meaningful patterns from enzyme-reaction pairs. (a) CDF of closest EC match between falsely predicted reaction and all true reactions of the query protein. MFP and RXNFP, left out for visualization purposes, are similar to DRFP and bag of molecules, respectively. (b) Closest EC match of false positive errors plotted as histograms. Colors are the same as in (a). Mean-max level is plotted as a vertical line for each model. (c) F1 scores on validation data are shown for RC aggregation models using varied numbers of message passings corresponding to varied bond-wise distances from the RC. Error bars are standard deviation. (d) Shows the similarity of two vectors in the learned embedding space. The first vector is the mean of all reaction embeddings sharing the same RC. The second is the embedding of the RC itself. Each pair of points corresponds to a unique RC featured in the dataset embedded either with the RC aggregation or bag of molecules model.

The particular number of bonds defining the constraint on RC aggregated’s context window arose from hyperparameter optimization. Performance is optimal in the model variant with four message passing steps (Fig. 6c). This model variant achieved the highest mean F1 and lowest variability over three independent data splits. Four message passing steps imply that the reaction embedding vector depends only on substrate features within three bonds of the RC. It is interesting to compare this radius to those used in the design of reaction templates for computer-aided biosynthesis workflows. RetroRules use a radius of 3-4 bonds, aligned with our result here^19,50^. Others generate templates of variable size through a heuristic, but they nevertheless focus on substrate features within the local neighborhood of the RC^51,52^.

To further elucidate RC aggregated’s reaction embeddings, we compared the mean of all reaction embeddings of a common RC with the embedding of the RC itself. The RC alone is in a cheminformatics sense a valid, balanced reaction which we can encode as a SMILES string and pass to our model. We expected that an encoder that had learned patterns relevant to enzymatic reactions would map reactions of a common RC near to the RC itself, each one differing slightly due to its specific characteristics. The mean of the full reaction vector embeddings should therefore be near the embedding of the RC itself. Using the RC aggregated encoder to embed reactions, we found that 94% of the time the full reactions’ vector mean had a similarity score of 0.8 or greater to the embedding of their shared RC. When using the bag of molecules to embed the reactions this dropped to 85%. Similarity scores were generated in the same manner as between a reaction and enzyme, which is also valid for this analysis since all inputs are mapped to the same vector space. Comparing RC by RC, the full reactions’ mean and RC embedding vector were closer in the majority (70%) of cases when encoded by the RC aggregated model compared to the bag of molecules model (Fig. 6d). This analysis is consistent with the idea that the RC aggregated model learns patterns that are more chemically meaningful.

## Discussion

In this study we developed and compared models capable of successfully predicting whether a specific enzyme catalyzes a specific reaction. The RC connected model exhibits consistently high performance across the variety of evaluation tasks in this study. Our models improved upon the performance of previous works despite the fact that they had much lower parameter counts and number of negative pairs sampled. Additionally, our maximum memory requirement is 650x lower than that of the closest benchmark, CLIPZyme, which utilizes *n* x *d* dimensional protein representations where we use *d* x 1, where *n* is the number of amino acid residues and *d* is the embedding dimension. This simplicity makes RC-GNN models well-suited for practical use without specialized hardware.

We demonstrated model generalization by extending methodologies routinely applied in protein property prediction tasks and multiclass classification. Following the example of prior work^24,38^, we used GSI to quantify pairwise similarity between enzymes. We extended this idea to pairwise reaction similarity with the RCMCS similarity metric. One can think of both GSI and RCMCS as methods that align graphs and score the alignments. In the case of amino acid sequences, graphs are linear. In the case of reactions, we have multiple disjoint subgraphs, the reactants and products. Maximum common subgraph with the extra condition to include the RC is an appropriate definition of alignment in this case. These similarity metrics served as inputs for our stratified similarity split method, which in turn enabled us to characterize model performance as a function of data point dissimilarity.

Like previous studies^24^, we observed that models performed extremely well over a wide range of enzyme dissimilarity. Estimates vary for the threshold on sequence identity after which protein function diverges. The most permissive is 40%^53^. Our results suggest all models we developed generalize well beyond the point at which memorizing sequence homology is effective. In part, we attribute this success to the representational power of protein language models^40^.

Generalization to dissimilar reactions is more difficult. The RC-GNN and CGR models performed best on this task. We attribute this to their ability to learn chemically meaningful patterns, revealed through false positive error analysis. We did not experiment with any form of semi-supervised technique for learning pre-trained reaction embeddings in contrast to the protein encoder. Future studies could investigate a form of masked language modeling appropriate for reaction graphs or related schemes.

The models developed here represent an important step forward in the formulation of enzymatic function prediction. Accommodating enzyme-reaction pairs as inputs removes ambiguity surrounding labels like EC number and multiplicity of reactions admitted by a given substrate. We share the belief that mapping reactions (especially previously unobserved reactions) to enzymes is a critical part of computer-aided biosynthesis^2,27,28^. Since our formulation of enzymatic function prediction is distinct from many previous studies, we took care to design a series of benchmarks, each to address a specific question. Building up from the least featured, the Morgan fingerprint model lacks a differentiable message passing function and thus reveals the benefit of learning these transformations from data as the GNNs do. The bag of molecules model adds differentiable message passing but leaves out RC information, equivalently an all-atom mapping of the reaction. Both the RC connected and CGR models make use of all-atom mapping in different ways but globally aggregate information in contrast to the more narrowly focused RC aggregated model.

A limitation of the RC aggregated, RC connected, and CGR models is the requirement of an all-atom mapping as an additional pre-processing step. There are tools to generate an all-atom mapping for a reaction including open-source and paid solutions^54^. One can also use minimal reaction templates to generate an all-atom mapping but are limited in this case to a set of known RCs^18,33^. Although all-atom mapping tools are not perfect, we show model prediction is robust to reported uncertainties in the atom mappings (Fig. S14). Moreover, the bag of molecules model performs well in many conditions and does not require an all-atom mapping. In addition to atom-mapping confidence, we provide extensive error analysis elucidating relative performance over EC subclasses, protein size, reactant and product size, and reactant and product number (Fig. S10-13).

There are several use cases for a model such as the one developed here. In the field of computer-aided biosynthesis, it can map highly dissimilar reactions generated by network expansion tools^55,56^ to amino acid sequences. It can be used to annotate amino acid sequences without direct experimental activity including those with functions inferred from homology, which we purposefully excluded from our dataset. We anticipate, like other state-of-the-art protein function annotation tools, our model is not a replacement for experiments^57^, but it is conveniently formulated to guide experiments and can therefore be used in an iterative loop of prediction and validation. In addition to biotechnology, we believe our model can aid hypothesis generation in fundamental studies of evolvability and robustness by identifying secondary enzymatic activities that can compensate for genetic perturbation in sometimes elaborate ways, such as has been described in the pyridoxal 5’-phosphate synthesis pathway in *E. coli*^4,6^.

## Methods

### Data curation and pre-processing

Enzyme-reaction pairs were collected from the UniProt and Rhea databases^31,32^. We pulled all manually annotated UniProt (SwissProt) entries containing information in the ‘Catalytic activity’ field. Using the Rhea IDs listed in the Catalytic activity field of each UniProt entry, we gathered reaction information in the form of reaction SMILES.

To standardize the reaction SMILES, we removed stereochemistry, neutralized charges that vary depending on environmental conditions (e.g., protonating-OH groups), applied standard molecule normalization transformations, and canonicalized molecular SMILES. Standardization was carried out with cheminformatics library rdkit^58^. All transport reactions (e.g., between cellular components) were removed. We automatically generated RCs for each reaction by applying a set of minimal enzymatic reaction operators^18^ to reactants, alternately protecting all but one substructure matching the operator’s RC template.

From enzyme entries we gathered amino acid sequences and subsequently input them into ESM-1b to generate vector embeddings. We excluded proteins listed as subunits of a multi-unit catalytic complex, and those with ‘Level of Evidence’ classified by UniProt as ‘Inferred from Homology’, ‘Predicted’, or ‘Uncertain’.

### Model implementation

Models were built using Chemprop^43^, Lightning and PyTorch. Reaction encoders are implemented as custom classes subclassed from one of the three libraries depending on the encoder. For the protein encoder, we used the pre-trained ESM-1b model^40^. ESM-1b embeddings were linearly transformed by a matrix whose weights were learned from the data. The prediction head consists of taking a dot product of the reaction and enzyme embeddings and then applying the sigmoid function to the result.

All model parameters were learned using a binary cross entropy loss function with a positive weight parameter chosen through hyperparameter optimization.

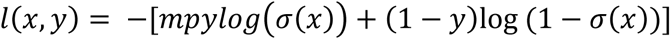

Where m = negative multiple, p = positive multiplier, y = label, x = model output.

CLIPZyme models were constructed with the hyperparameters found optimal in Mikhael et al., 2024, including a linear warmup cosine learning rate schedule^27^. For all experiments but the time-based split, a model with the CLIPZyme pseudo-transition-state reaction encoder and linear ESM-1b protein encoder were trained on the data set curated in this work. For the time-based split, we queried the model trained in Mikhael et al., 2024 which was provided with the published code and data.

We trained a ReactZyme model with ESM2^59^ protein embeddings and Uni-Mol^60^-based reaction embeddings as this architecture achieved top performance in most of the evaluations in Hua at al., 2024^28^. In lieu of a provided trained model, we reproduced their training procedure with the accompanying code and data.

### Data splitting and negative sampling

In order to systematically evaluate model generalization error, we used a data splitting approach that we call stratified similarity split. Like normal stratified splitting, we arrange data across splits in a way that preserves representation of multiple categories. In stratified similarity split, a category is characterized by a range of maximum cross-split similarity score. For example, a data point assigned to a validation fold in the [0.4-0.6) category is between 0.4 and 0.6 similar to the nearest training fold data point. Stratified similarity split is applied in both the outer (test / training-validation) and inner splits (training / validation).

Stratified similarity split is achieved by first clustering the dataset according to a similarity metric up to a chosen upper bound on similarity, sampling clusters, relaxing the bound to the next value, and repeating the process up to the most permissive similarity bound. We used both protein-based and reaction-based similarity metrics for splitting and evaluation in this work. On the protein side, amino acid sequences were locally aligned and clustered using the algorithm cd-hit^61^. Similarity between pairs of sequences is defined as the ratio of matching residues to the number of residues in the shorter of the two sequences. On the reaction side, we sought an analogous method of computing pairwise similarity based on aligning graphs, which can be branched and/or cyclical in the case of reactions, in contrast to amino acid sequences. We therefore used rdkit^58^ to find the maximum common subgraph (MCS) between reaction graphs. We further required that the RC be contained in the MCS to ensure that the similarity metric would be sensitive to structural features relevant for the reaction and not spurious, non-instructive shared structural features. We call the custom MCS procedure RCMCS. To compute a bounded similarity score, we take the ratio of atoms in the RCMCS to the number of atoms in the larger of the two reaction graphs. A similarity score of 1.0 occurs exactly when the two reactions are identical. Example pairs of RCMCS-compared reactions with other values are given in Figure 3b.

In addition to the known positive enzyme-reaction pairs which we sourced from UniProt and Rhea, we assumed a number of negative pairs by either global selection or ARC-based generation of unobserved enzyme-reaction pairs from our dataset. We sampled negatives in a ten-to-one ratio with our positive pairs in test and validation folds (unless otherwise stated) and in a three-to-one ratio in training folds. The latter ratio was determined during hyperparameter optimization.

ARC negative sampling utilizes the fact that there are sometimes multiple moieties on reactants identical to the reaction center^37^. These “alternate reaction centers” are however not acted on by the transformation of the observed reaction. We infer from this that the structures surrounding these alternate reaction centers somehow preclude the necessary catalytic activity of the enzyme. In other words, the reactions that would have resulted can be considered negative pairs of the enzyme in question. We generate all ARC reactions by applying the minimal rule^18^ corresponding to each observed reaction to all RCs other that the one transformed in the reaction of interest. We then cross reference the ARC reactions with our existing enzyme-reaction adjacency matrix to form negative pairs.

For the time-based split experiment, we downloaded reactions from Rhea and enzyme sequences from UniProt on September 15, 2025. We subsequently removed all reactions and proteins occurring in either our data set (downloaded March 10, 2024) or the data sets used in Mikhael et al., 2024 and Hua et al., 2024. We globally selected negative pairs in a 10:1 ratio to align with the negative sampling method used in both these prior works.

### Hyperparameter optimization

We selected values for hyperparameters using Bayesian methods, specifically the Tree-structured Parzen Estimator and Hyperband Pruner, implemented in the Optuna package^62^. We maximized ROC-AUC scores on validation data. Number of training epochs were determined with retrospective optimal checkpoint selection on validation data. Each baseline model independently underwent this procedure. A full list of optimized hyperparameters and their values appear in Table S1.

## ASSOCIATED CONTENT

### Supporting Information

Additional details and validation of the dataset used, the method of stratified similarity splitting, and optimal hyperparameters selected through cross validation.

## AUTHOR INFORMATION

### Corresponding Author

Keith E.J. Tyo

Telephone: +1 847 868 0319

Fax: +1 847 491 3728

### Author Contributions

The manuscript was written through contributions of all authors. All authors have given approval to the final version of the manuscript.

## ACKNOWLEDGMENT

This work was performed as part of the Bio-Optimized Technologies to keep Thermoplastics out of Landfills and the Environment (BOTTLE) Consortium and was supported by AMO and BETO under contract no. DE-AC36-08GO28308 with the National Renewable Energy Laboratory (NREL), operated by Alliance for Sustainable Energy, LLC.

Stefan Pate was supported in part by the National Institutes of Health Training Grant (T32GM153505 and T32GM008449) through Northwestern University’s Biotechnology Training Program.

We would like to thank Yash Chainani, Geoffrey Bonnanzio, and Pavel Katsev for helpful discussion.

This research was supported in part through the computational resources and staff contributions provided for the Quest high performance computing facility at Northwestern University which is jointly supported by the Office of the Provost, the Office for Research, and Northwestern University Information Technology.

## DATA AND SOFTWARE AVAILABILITY

All code and data associated with this work are made available at: github.com/stefanpate/rcgnn

