## Supplementary Information for "RC-GNN: A predictive model of enzyme-reaction pairs"

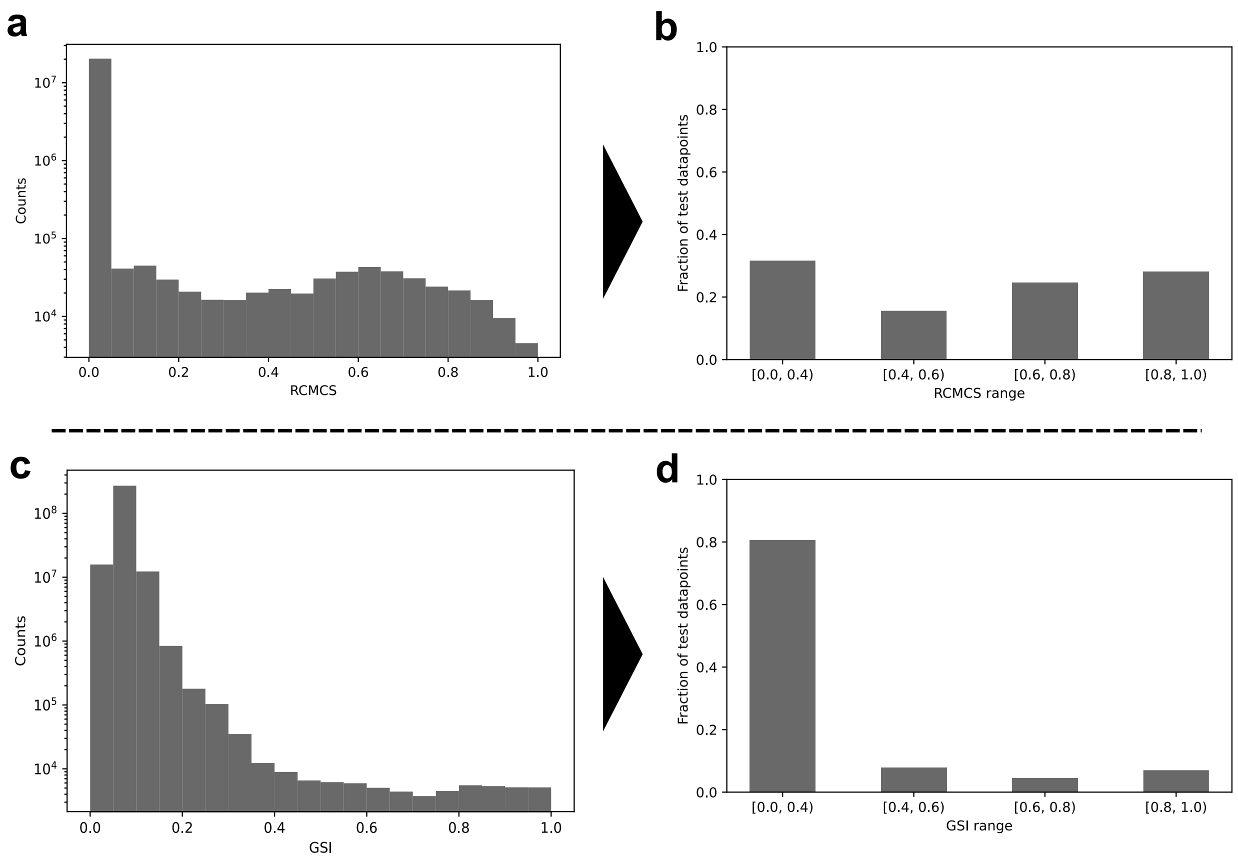


**Figure S1** Stratified similarity split ensures the maximum similarity between test and train data points varies widely. (a) Histogram showing the background distribution of pairwise reaction similarities over the whole data set. (b) Stratified similarity split generates a test set approximately balanced in representation of similarity-defined subsets. (c) Same as (a) for pairwise protein similarity. (d) Stratified similarity split guarantees representation of similarity-defined subsets but is limited by the underlying distribution of pairwise similarities.

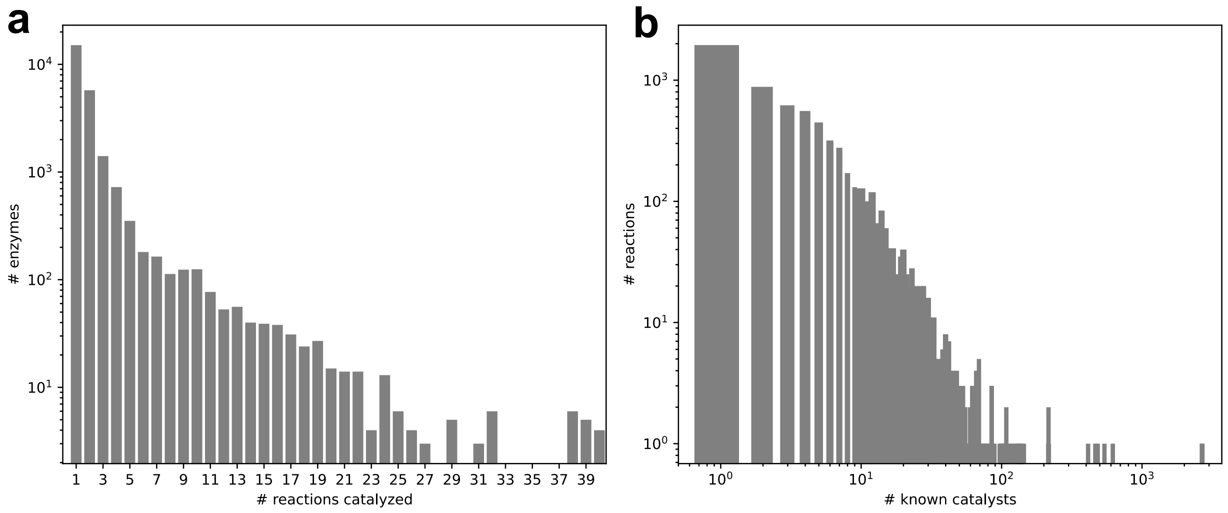


**Figure S2** Dataset includes enzymes catalyzing multiple reactions and reactions catalyzed by multiple enzymes. (a) Histogram showing counts of unique enzymes catalyzing a given reaction (2.0 ± 2.6). (b) Histogram showing counts of unique reactions catalyzed by a given enzyme (7.6 ± 50.4)

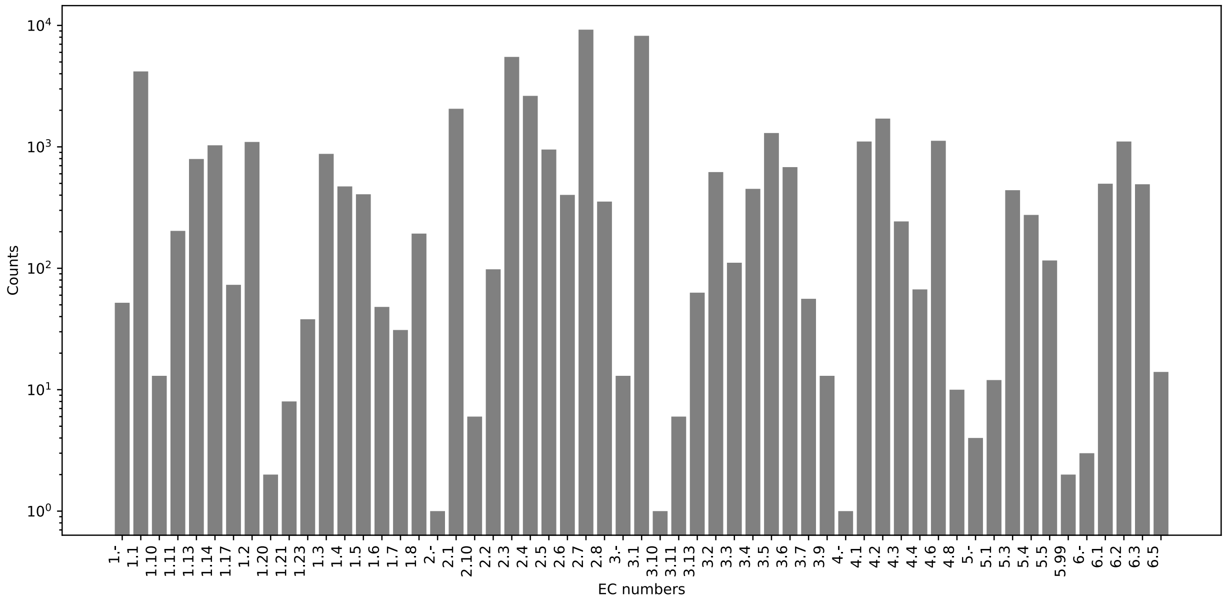


**Figure S3** Dataset covers six major top-level EC numbers. Transport reactions were not included.

**Table S1** Optimal hyperparameters used for each model architecture

| | RC aggregated | RC connected | Bag of molecules | CGR | RXNFP | DRFP | Morgan fingerprint |
|---|---|---|---|---|---|---|---|
| Training epochs (RCMCS/ Reaction/ Reaction Center) | 12/21/6 | 9/24/12 | 9/24/24 | 12/15/24 | 12/9/24 | 15/21/9 | 12/24/3 |
| Negative multiple | 4 | 4 | 4 | 6 | 3 | 3 | 3 |
| Positive multiplier | 1 | 1 | 1 | 2 | 1 | 3 | 3 |
| Reaction encoder type | GNN | GNN | GNN | GNN | Linear | MLP | MLP |
| Embedding dimension | 145 | 48 | 24 | 273 | 55 | 25 | 56 |
| # message passings | 4 | 4 | 5 | 6 | n/a | n/a | n/a |
| # reaction encoder layers | n/a | n/a | n/a | n/a | n/a | 4 | 4 |
| ECFP radius | n/a | n/a | n/a | n/a | n/a | 3 | 2 |
| Input vector length | n/a | n/a | n/a | n/a | 256 | 2048 | 2048 |

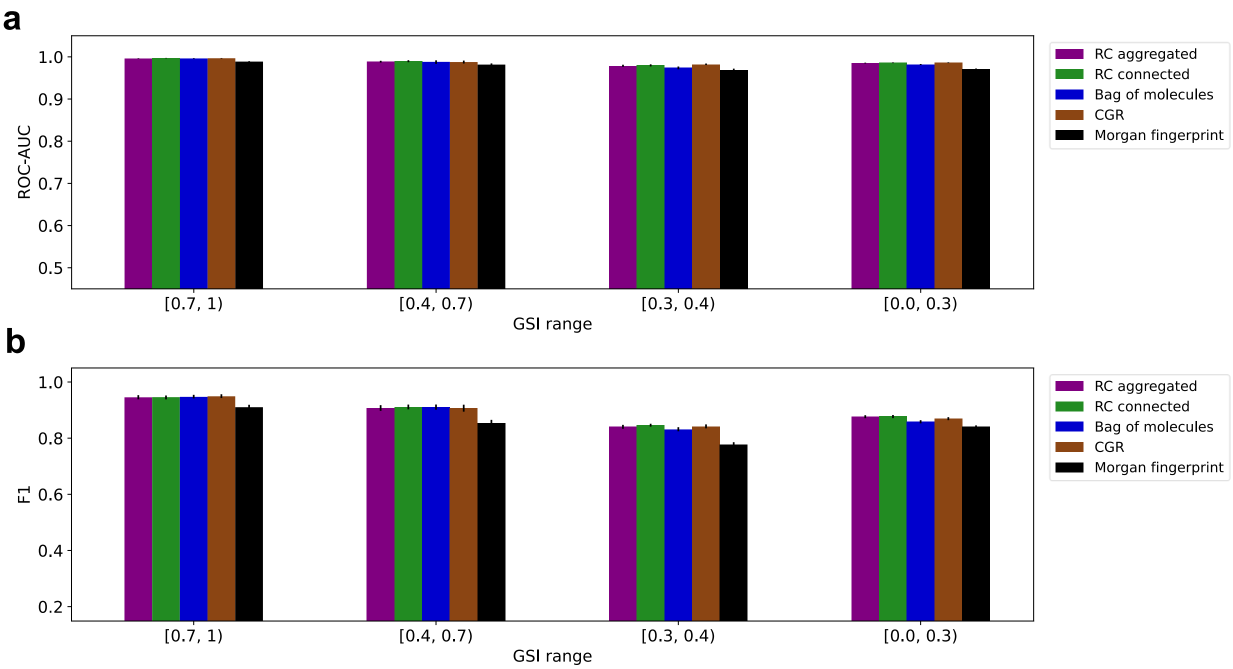


**Figure S4** Model performance split by max GSI between test and train points. (a) ROC-AUC. (b) F1 score. Error bars are 95% confidence intervals computed by bootstrapping (n=20).

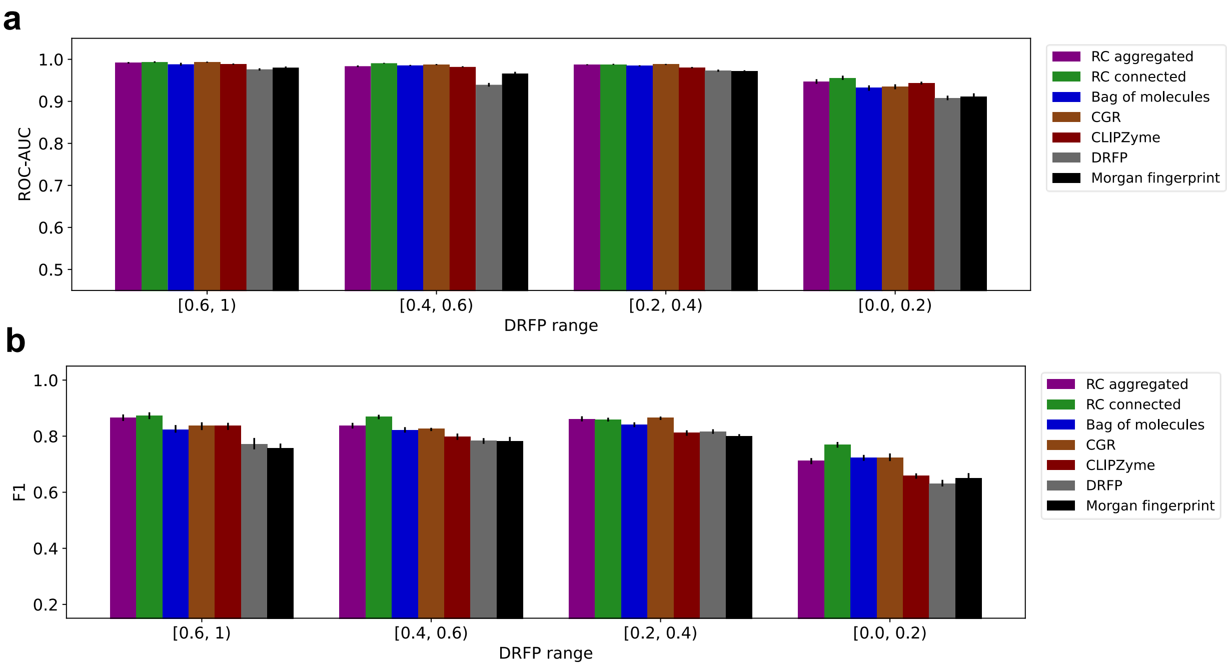

**Figure S5** Model performance split by max Tanimoto similarity of test and train DRFP representations. (a) ROC-AUC. (b) F1 score. Error bars are 95% confidence intervals computed by bootstrapping (n=20).

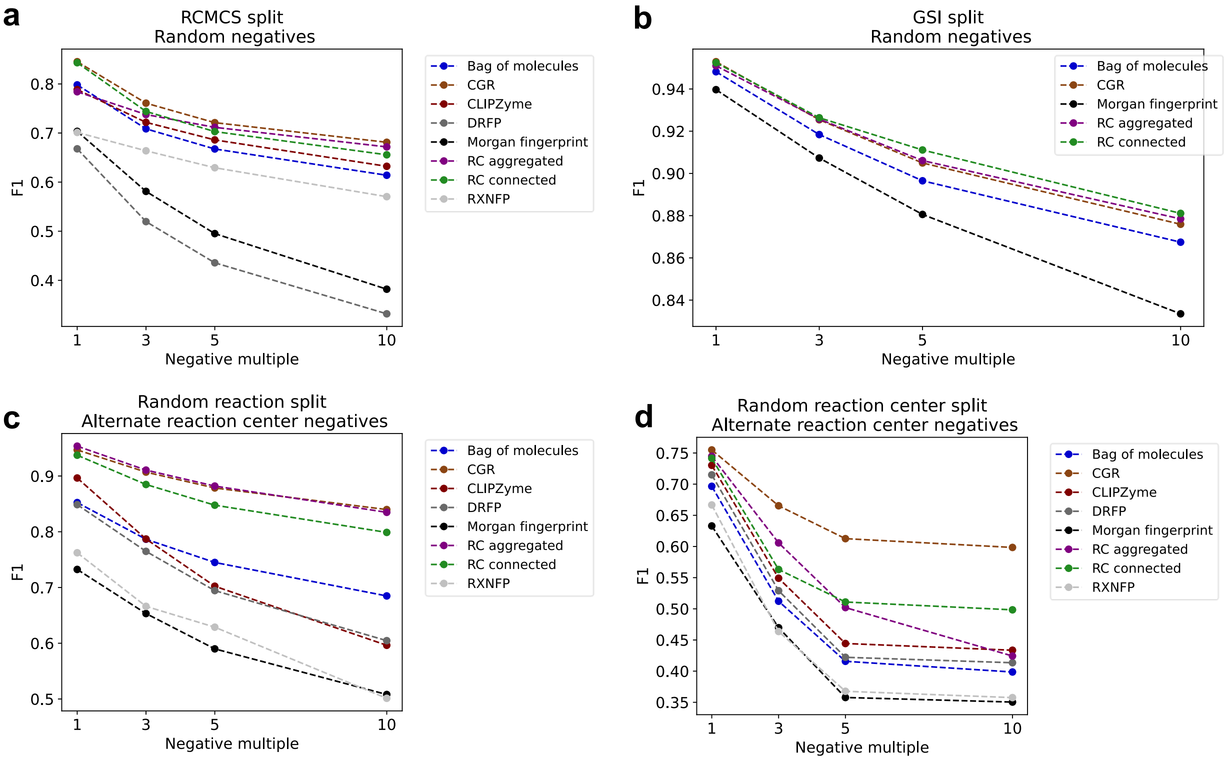


**Figure S6** F1 scores as a function of the ratio between negative and positive sample in the test set. (a) RCMCS split and random negative sampling. (b) GSI split and random negative sampling. (c) Random reaction split and ARC negative sampling. (d) Random reaction center split and ARC negative sampling.

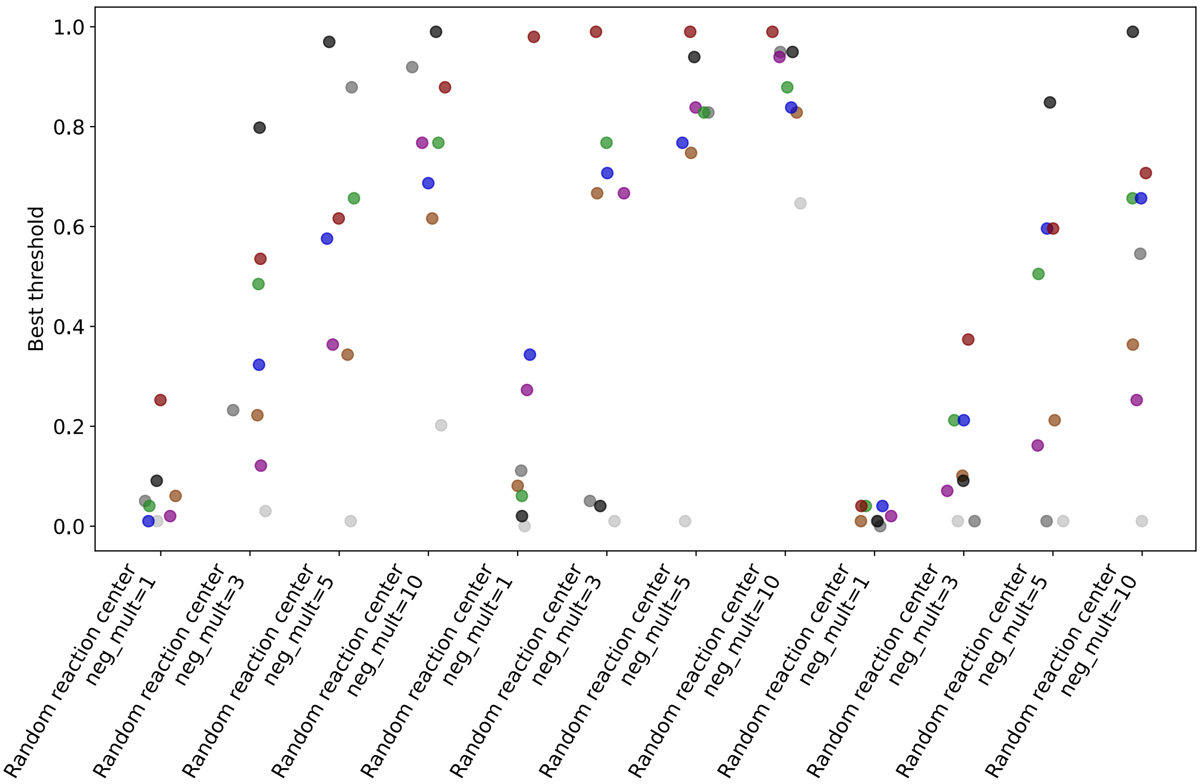


**Figure S7** F1-maximizing decision threshold for all combinations of data splitting strategy, negative multiple, and model architecture. Model architecture is denoted by same color scheme used throughout the manuscript: RC aggregated = purple, RC connected = green, Bag of molecules = blue, Morgan FP = black, CLIPZyme = dark red, CGR = dark orange, DRFP = dark gray, RXNFP = light gray.

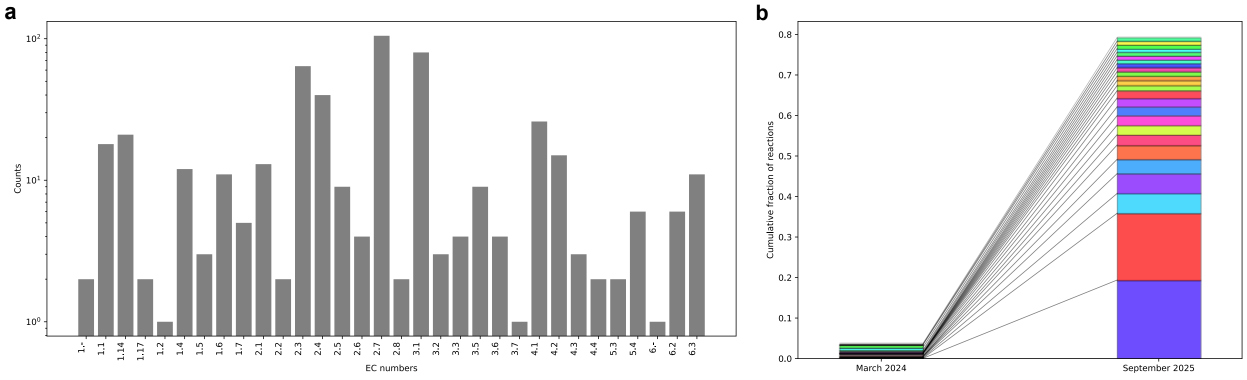


**Figure S8** Characterization of time split experiment. (a) Time split reactions cover six major top-level EC numbers. Transport reactions were not included. (b) Time split reactions have reaction centers which are relatively rare in the training data set.

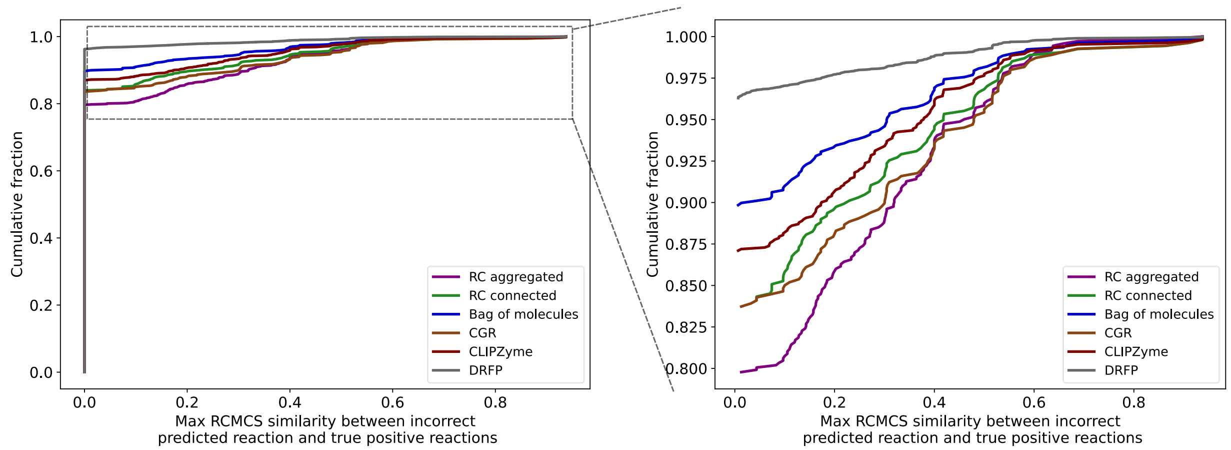


**Figure S9** RC aggregated false positive errors tend to be the most chemically reasonable. (Left) CDF of maximum RCMCS similarity between the incorrectly predicted reaction and reactions observed to be catalyzed by the query protein. (Right) Magnification of non-zero range of RCMCS similarity.

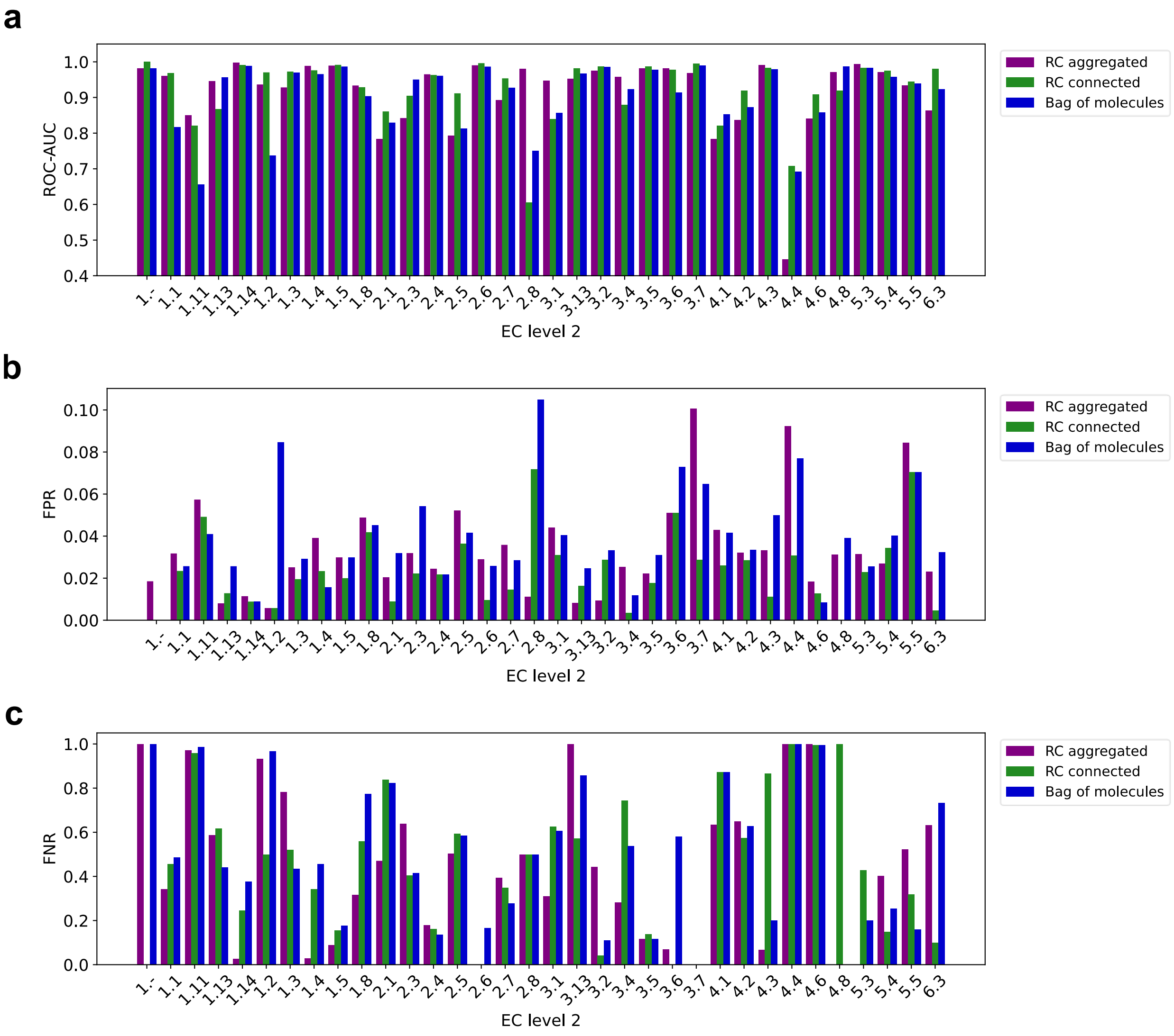


**Figure S10** Error analysis for EC level 2 number. (a) ROC-AUC (b) False positive rate (c) False negative rate

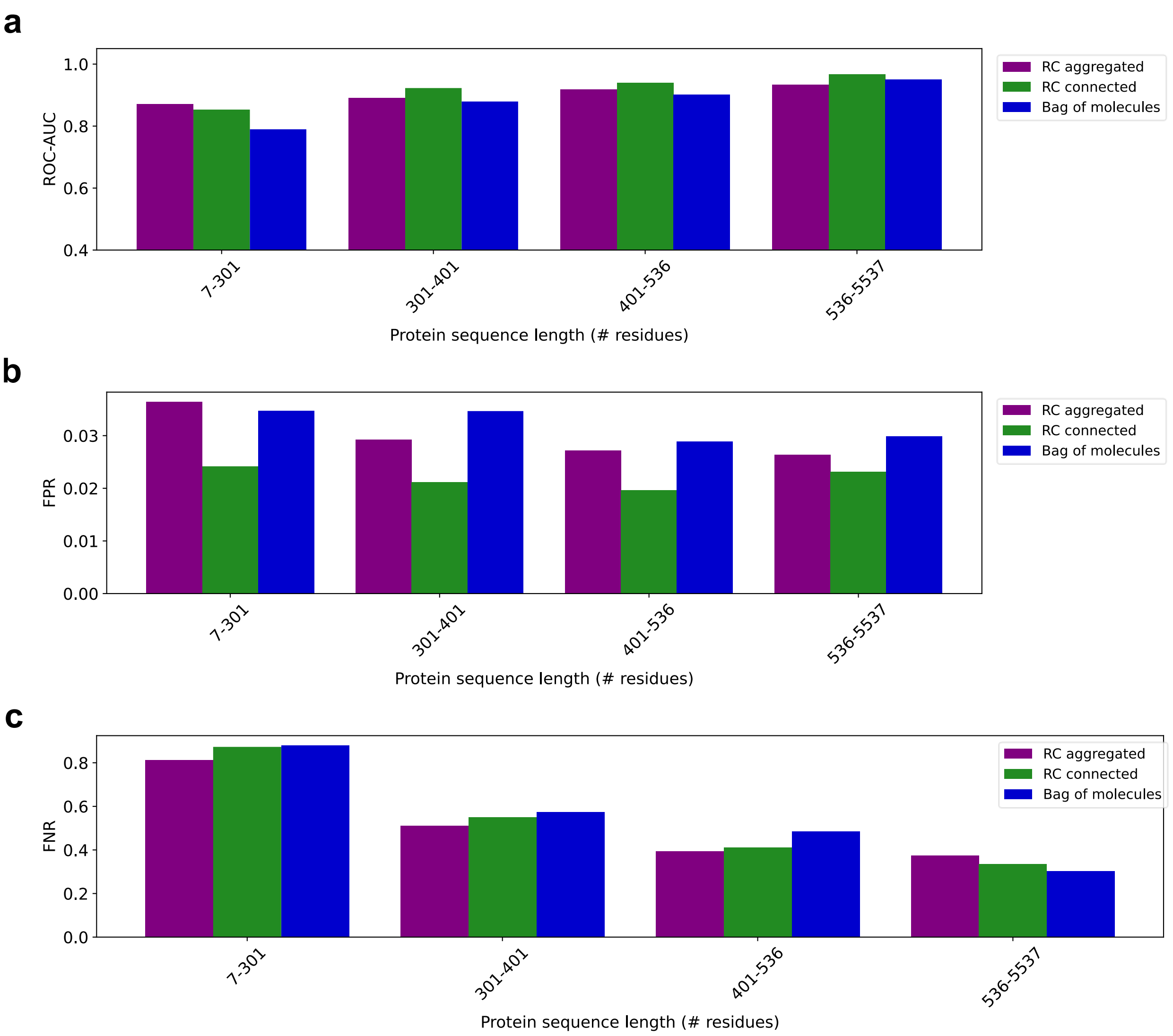


**Figure S11** Error analysis for protein sequence length. (a) ROC-AUC (b) False positive rate (c) False negative rate

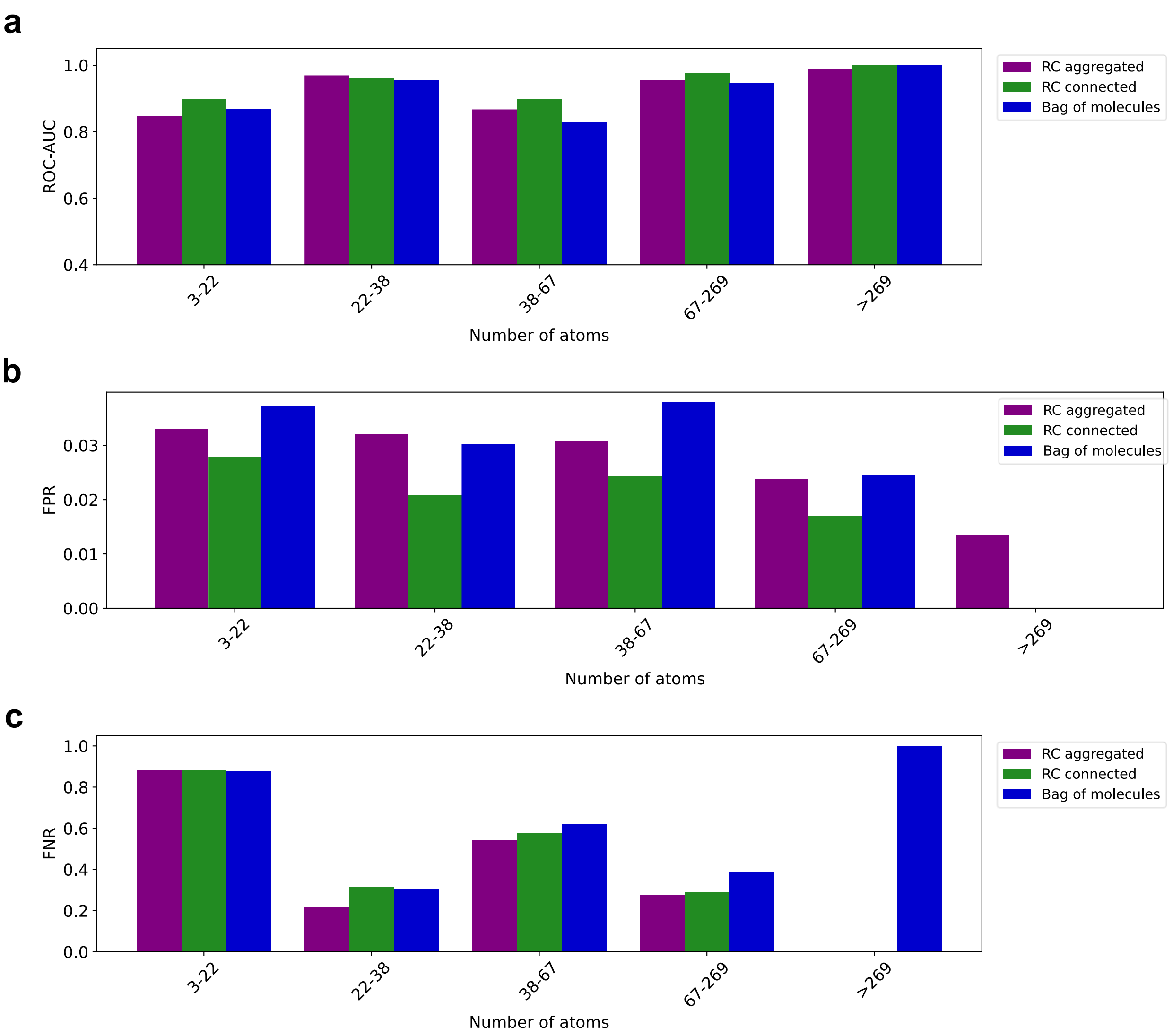


**Figure S12** Error analysis for total number of atoms among reactants and products. (a) ROC-AUC (b) False positive rate (c) False negative rate

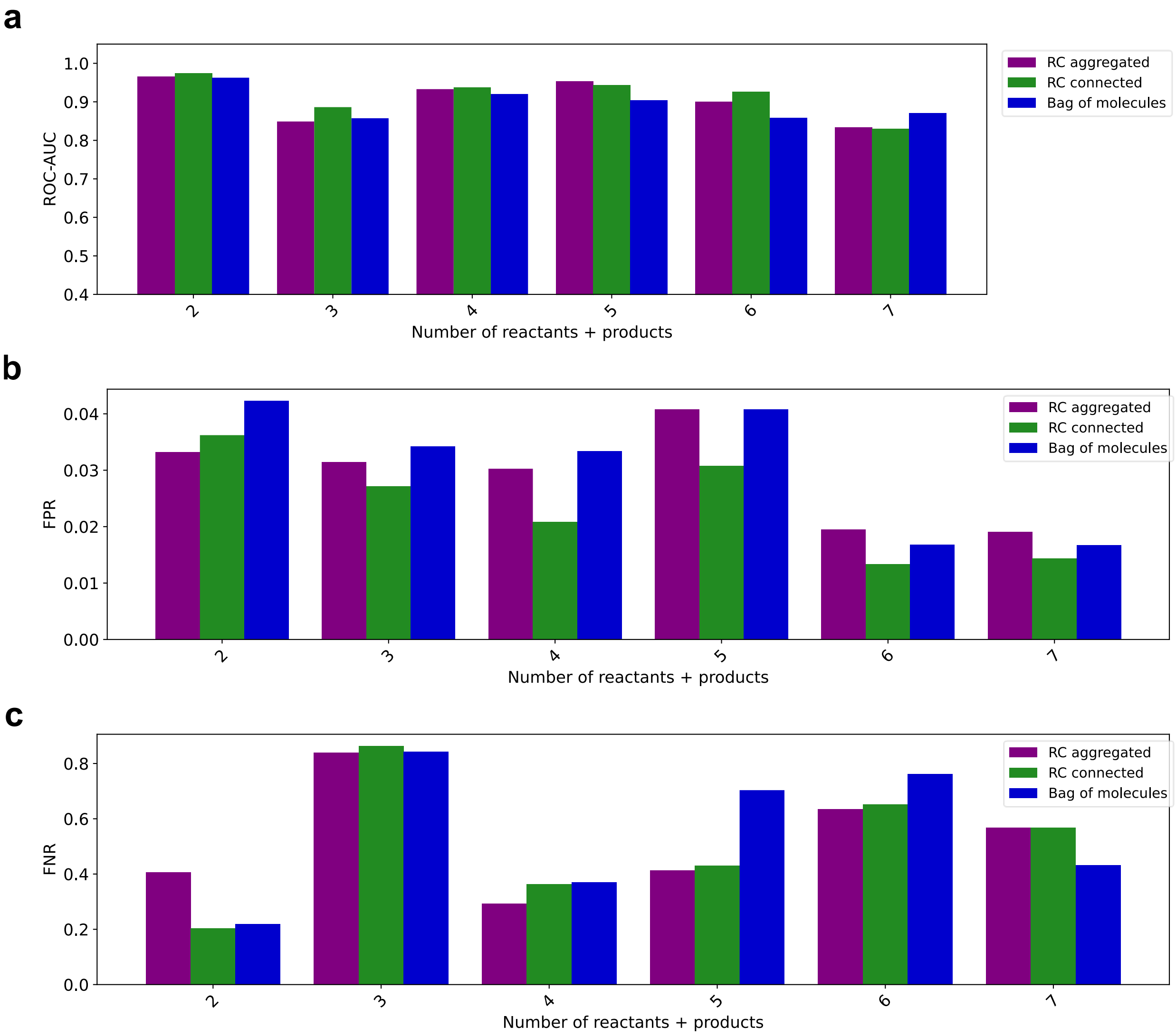


**Figure S13** Error analysis for number of reactant and product molecules involved in the reaction. (a) ROC-AUC (b) False positive rate (c) False negative rate

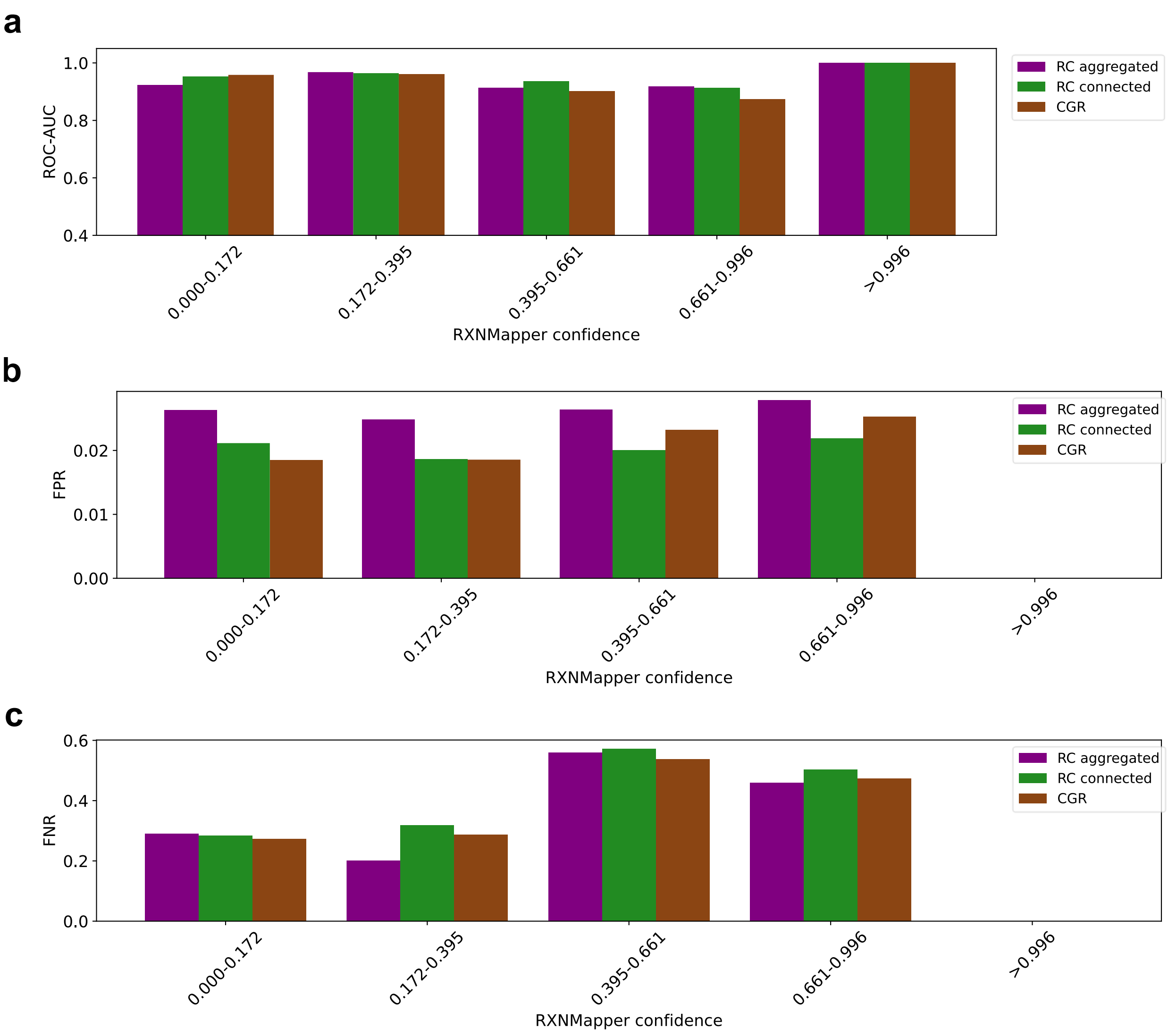


**Figure S14** Error analysis for RXNMapper (all-atom-mapping tool) confidence score. All-atom-mapping-dependent models are robust to variations in confidence. (a) ROC-AUC (b) False positive rate (c) False negative rate
